# GraphUnzip: unzipping assembly graphs with long reads and Hi-C

**DOI:** 10.1101/2021.01.29.428779

**Authors:** Roland Faure, Nadège Guiglielmoni, Jean-François Flot

## Abstract

Long reads and Hi-C have revolutionized the field of genome assembly as they have made highly continuous assemblies accessible for challenging genomes. As haploid chromosome-level assemblies are now commonly achieved for all types of organisms, phasing assemblies has become the new frontier for genome reconstruction. Several tools have already been released using long reads and/or Hi-C to phase assemblies, but they all start from a linear sequence, and are ill-suited for non-model organisms with high levels of heterozygosity. We present GraphUnzip, a fast, memory-efficient and accurate tool to unzip assembly graphs into their constituent haplotypes using long reads and/or Hi-C data. As GraphUnzip only connects sequences in the assembly graph that already had a potential link based on overlaps, it yields high-quality gap-less supercontigs. To demonstrate the efficiency of GraphUnzip, we tested it on a simulated diploid *Escherichia coli* genome, and on two real datasets for the genomes of the rotifer *Adineta vaga* and the potato *Solanum tuberosum*. In all cases, GraphUnzip yielded highly continuous phased assemblies.

## 1 Introduction

The field of genomics is thriving and chromosome-level assemblies are now commonly achieved for all types of organisms, thanks to the combined improvements of sequencing and assembly methods. Chromosome-level assemblies are generally haploid, regardless of the ploidy of the genome. To obtain a haploid assembly of a multiploid (i.e. diploid or polyploid) genome, homologuous chromosomes are collapsed into one sequence. However, assemblers often struggle to collapse highly heterozygous regions, which leads to breaks in the assembly and duplicated regions [1]. Furthermore, haploid assemblies provide a partial representation of multiploid genomes; ideally, multiploid genomes should be phased rather than collapsed if the aim is to grasp their whole complexity [2].

The combination of low-accuracy long reads, such as Oxford Nanopore Technologies reads (ONT) and Pacific Biosciences (PacBio) Continuous Long Reads (CLR), and proximity ligation (Hi-C) reads has made chromosome-level assemblies accessible for all types of organisms. The latest development of PacBio, high-accuracy long circular consensus sequencing (CCS) reads (a.k.a. HiFi), is now starting to deliver highly continuous phased assemblies [3, 4, 5]. Existing tools are able to use either long reads (Falcon-Unzip [6], WhatsHap [7]) or Hi-C reads (Falcon-Phase [8], ALLHiC [9]) for phasing assemblies, but they are limited to phasing local variants or well-identified haplotypes and are not suited for complex, highly heterozygous genomes.

We present GraphUnzip, a new tool to phase assemblies using long reads and/or Hi-C. GraphUnzip implements a radically new approach to phasing that starts from an assembly graph instead of a collapsed linear sequence. In an assembly graph, heterozygous regions result in bubbles every time the assembler is unable to collapse the haplotypes or to choose one of them. GraphUnzip “unzips” the graph, meaning that it separates the haplotypes by: 1) duplicating homozygous regions that have been collapsed; 2) partitioning heterozygous regions into haplotypes. As it takes as input and produces as output an assembly graph, our tool only connects contigs that are actually adjacent in the genome and yields gap-less scaffolds, i.e. supercontigs. We tested GraphUnzip on a simulated diploid *Escherichia coli* genome, and on the genomes of the rotifier *Adineta vaga* and the potato *Solanum tuberosum*. Graphunzip is available at github.com/nadegeguiglielmoni/GraphUnzip.

## 2 Methods

### 2.1 Inputs

GraphUnzip requires an assembly graph in GFA format. The Hi-C input is a sparse matrix, such as the one obtained when processing the reads with hicstuff [10]. The long reads are mapped to the assembly graph using GraphAligner [11].

### 2.2 Overview of GraphUnzip

In an assembly graph, contigs (segments) that are inferred to be adjacent or overlap in the assembly are connected with edges. However, some of these connections between contigs may be artefacts. To discriminate correct links from erroneous ones, GraphUnzip relies on long reads and/or Hi-C data. These data are translated to interactions between segments: two segments have a strong interaction based on long reads when many reads align on both segments; strong Hi-C interactions correspond to frequent Hi-C contacts between the two segments.

GraphUnzip first builds one or two interaction matrices, depending on whether long-read data, Hi-C data or both are provided (Figure 1). GraphUnzip reviews all segments and their links. For each link, an interaction intensity value *i* is computed based on long reads data; if no link can be categorically deleted based on this data, the intensity value is computed based on Hi-C data.

**Figure 1:**
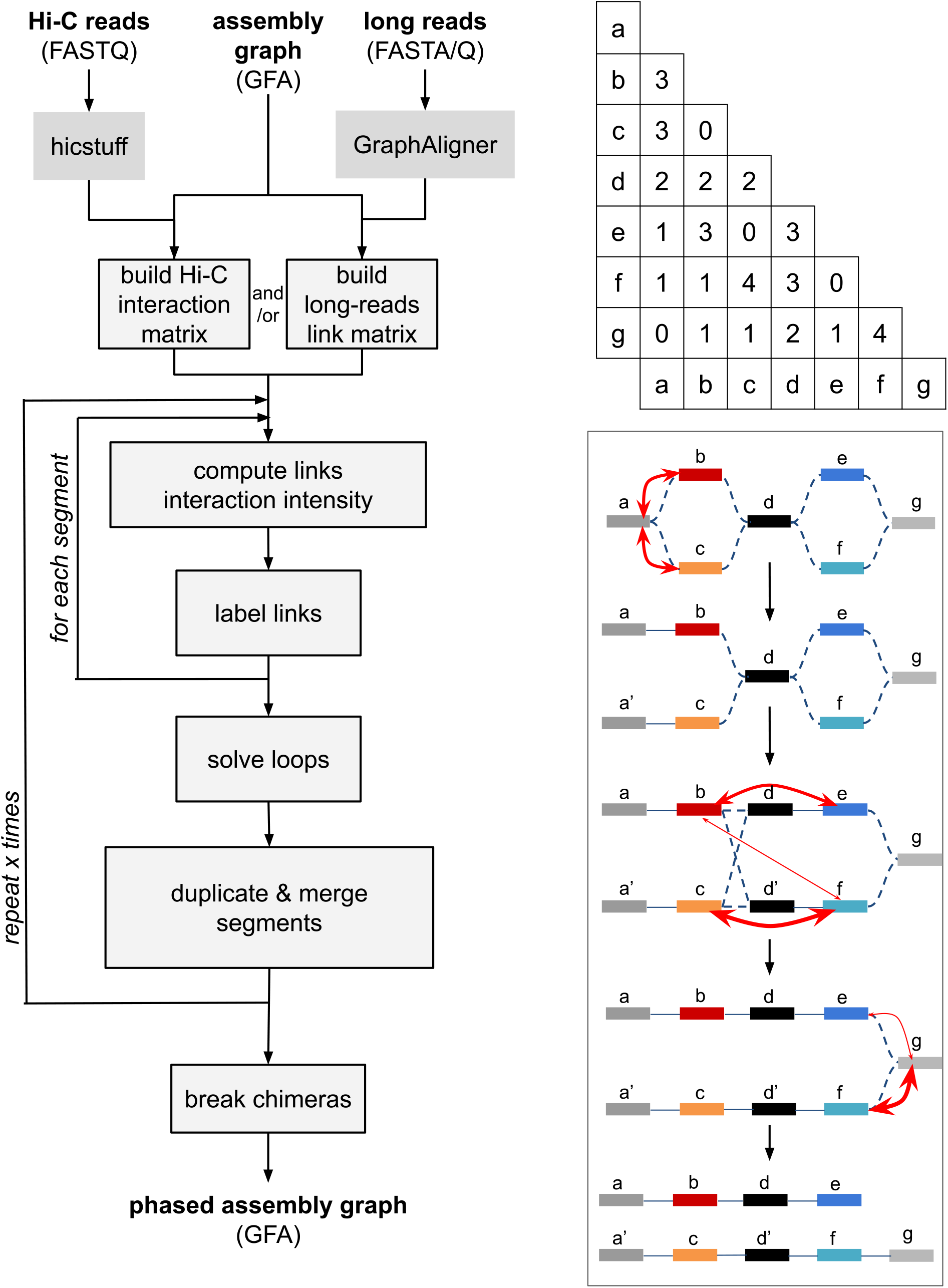
Description of GraphUnzip: workflow of the program (left), interaction matrix (top right), and overview of the algorithm to discriminate links (bottom right). This example algorithm analyzes the potential links between the segments a, b, c, d, e, f, g. The red arrows represent the intensity of interactions between the segments, computed based on the values in the matrix.

When assessing two putative links A-B and A-C between segments A, B and C, the respective strengths of these links are calculated as the number of contacts (long reads or Hi-C) exclusive to A and B vs. the number of contacts exclusive to A and C. For example, in the third step of Figure 1, when trying to associate segment (a,b) to either (d,e) or (d’,f), (a,b) shares its contacts with contig d/d’ between all its neighbors, so only the contacts with e and f are considered.

When one segment has several potential links to other segments, these links are compared in a pairwise fashion. This comparison is made using two user-provided thresholds: the rejection threshold *T_R_* and the acceptance threshold *T_A_*, where *T_R_* < *T_A_*. Considering two links X and Y and their respective interaction values *i*(*X*) and *i*(*Y*), if *i*(*X*) < *i*(*Y*): if *i*(*X*)/*i*(*Y*) < *T_R_*, then the link X is removed; else, if *i*(*X*)/*i*(*Y*) < *T_A_*, the link is flagged as dubious, as GraphUnzip is unable to make a decision. The link is considered correct when *i*(*X*)/*i*(*Y*) ≥ *T_A_*. Then, if long reads data is provided, GraphUnzip trims loops (where one segment has a potential link to itself or to both ends of another segment). Weak links are removed and every segment that has several correct links is duplicated. These segments are typically collapsed homozygous regions that need to be duplicated to be phased with each allele. Every copy of the duplicated segment keeps the links of the original segment at its other end. This entails that the duplication of segments creates many new links.

The links are iteratively processed to entirely phase the assembly for *s* steps, where *s* is a user-provided parameter. Because extremely long segments tend to share a significant number of Hi-C contacts even if they are not adjacent, we observed that in extreme cases the algorithm could join two chromosomes by their telomeric ends. The Hi-C matrix is used at the end of the process to detect such chimeric connections in the assembly graph, based on low Hi-C interactions, and break them.

### 2.3 *Escherichia coli* simulations and assembly

We simulated a diploid *Escherichia coli* by randomly mutating 200 blocks of 10 kilobases (kb) with a 3% error rate. Based on this genome, we simulated 250 basepairs (bp) paired-end reads with a 1% error rate (as is usual for Illumina reads), a 100X coverage, using the tool sim_reads from the package IDBA [12].

These reads were assembled using Bwise [13], available at github.com/Malfoy/Bwise. Bwise is a de Bruijn assembler that takes as input short reads with a coverage of 50X or more, and provides as output both the linear sequence in FASTA and the assembly graph in GFA format. It does not collapse heterozygous regions, making it appropriate for creating an interesting assembly to phase. The output was a typical partly diploid, partly haploid assembly, with a total assembly length of 8.3 Mb. In GFA format, links have an ‘overlap’, meaning that a few nucleotides at the end of the first contig are also present at the beginning of the other. As long overlaps can lead to many multimapped Hi-C reads, we modified a version of gimbricate (github.col/ekg/gimbricate) and used it to recompute the graph without overlaps.

With sim3C (github.com/cerebis/sim3C) [14], we simulated 1 million 150 bp paired-end Hi-C reads from the complete mock genome, based on a DpnII preparation. The Hi-C reads were mapped and preprocessed with hicstuff, with 98% mapping rate. GraphUnzip was run with parameters --`accept 0.20` --`reject 0.10`. The dotplot (Figure 2) was obtained by mapping the supercontigs longer than 100 kb, outputted by GraphUnzip, against the mock genome with minimap2 [15] with the parameter -`x asm5` and then the plot was built with minidot (github.com/lh3/miniasm) with parameter -`i 1.00`.

**Figure 2:**
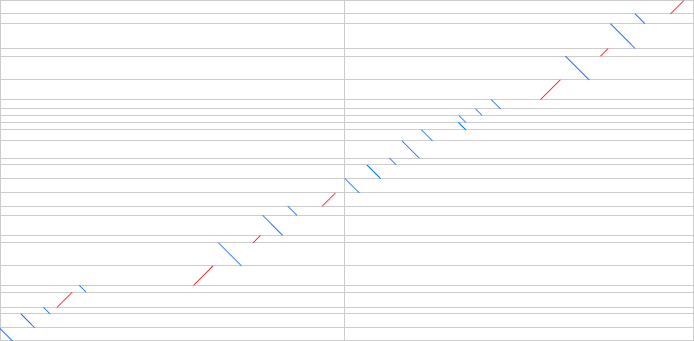
Comparison of the GraphUnzip supercontigs longer than 100 kb (y axis) versus the two chromosomes of the reference simulated diploid genome of *Escherichia coli* (x axis).

### 2.4 *Adineta vaga* assemblies

HiFi reads were generated at the Leiden Genome Technology Center. The HiFi dataset had a total size of 30.8 Gb and a N50 = 16.9 kb. Other reads were published in [16], and consist of: ONT (17.5 Gb, N50 = 18.8 kb), PacBio CLR (23.5 Gb, N50 = 11.6 kb), Illumina (2*250 bp, 11.4 Gb), Hi-C (2*66 bp, 55 million pairs). The full HiFi dataset was assembled with Flye [3] with the parameters --`pacbio-hifi reads` --`keep-haplotypes`. HiFi reads longer than 20 kb were assembled with hifiasm [4] with the parameter −l 0, and the p_utg assembly graph was used. Assembling the full dataset with hifiasm would yield an oversized assembly (258 Mb without overlaps). Low-accuracy long reads (ONT or PacBio CLR) were corrected with the Illumina reads using Ratatosk [17] to generate high-accuracy long reads, with default parameters. These corrected long reads were assembled with Flye using the same parameters as for HiFi assemblies. Long reads were mapped to the assembly graph using GraphAligner with the parameter -`x vg`. Hi-C reads were processed with hicstuff using the parameters --`aligner bowtie2` --`enzyme DpnII` --`iterative`. For all runs of GraphUnzip, the parameters were set to --`exhaustive` --`whole_match --minimum_match 0.8` and only --`accept` and --`reject` were adapted: Flye HiFi assembly --`accept 0.30 --reject 0.10`; hifiasm HiFi assembly --`accept 0.25 --reject 0.10`; corrected ONT assemblies --`accept 0.30 --reject 0.30`; corrected PacBio CLR assemblies --`accept 0.50 --reject 0.20`. The GFA was then analyzed with Bandage [18], available at rrwick.github.io/Bandage/, and all non-ambiguous paths were merged. The Flye and hifiasm assemblies of HiFi reads were also scaffolded with ALLHiC to compare with the GraphUnzip supercontigs. First, the Hi-C reads were mapped to the draft assemblies using the Burrows-Wheeler Aligner [19] with `bwa aln` and bwa sampe. Then the mapped reads were processed with the scripts `PreprocessSAMs.pl` (with parameter `GATC`) and `filterBAM_forHiC.pl`. Using the processed mapped Hi-C reads, the contigs of the draft assembly were partitioned in 12 groups with `ALLHiC_partition -k 12`. The data and groups were further processed with `allhic extract --RE GATC` and `allhic optimize`, and finally the contigs were scaffolded using `ALLHiC_build`.

### 2.5 *Solanum tuberosum* assemblies

Reads published in [20] were retrieved from the NCBI Sequence Read Archive with the Bioproject accession number `PRJNA573826`. The HiFi reads were assembled with hifiasm with the parameter -`l 0`, and we then used the p_utg assembly graph. All the HiFi reads and the ONT reads longer than 25 kb were mapped to the assembly using GraphAligner with the parameter -`x vg`. Hi-C reads were processed with hicstuff using the parameters --`aligner bowtie2 --enzyme MboI --iterative`. GraphUnzip was run with parameters --`accept 0.40 --reject 0.10 --exhaustive --whole_match --minimum_match 0.8`. All non-ambiguous paths in the GFA were merged with Bandage.

### 2.6 Assemblies evaluation

We used calN50 available at github.com/lh3/calN50 to compute NG50s. The NG50 was computed against an estimated size of 192.6 Mb for *Adineta vaga*, as the haploid genome size was estimated to 96.3 Mb [16], and 1.67 Gb for *Solanum tuberosum* (published assembly size [20]). BUSCO v4 [21] was run with parameters -`m genome --long` against the dataset metazoa odb10 for *Adineta vaga*, viridiplantae odb10 for *Solanum tuberosum*. The contact maps of *Adineta vaga* were built using the hicstuff pipeline, as described previously, and with the commmand `hicstuff view --binning 30`.

### 2.7 Computational performance

RAM usage and CPU time were measured with the command `/usr/bin/time -v`. *Adineta vaga* was tested on a laptop with 16 GB of RAM and a i7-8550U 1.8 GHz processor. *Solanum tuberosum* was tested on a desktop computer with 128 GB of RAM and a i9-9900X 3.5 GHz processor.

## 3 Results

### 3.1 Simulated data

To check that GraphUnzip behaved as expected, we first tested it on a simulated diploid *Escherichia coli* genome (O157:H7 Sakai strain). To obtain a diploid organism, the whole genome was duplicated and we created an alternation of conserved and mutated regions along the genome, as would be expected in a real diploid organism. The final simulated genome reached a size of 11.2 Megabases (Mb) and a heterozygosity rate of 1%. The draft assembly of simulated high-accuracy short reads reached a size of 8.3 Mb. We unzipped the assembly graph using simulated Hi-C reads, and the assembly size grew to 10.7 Mb, closer to the expected size. The NG50 rose from 10 to 107 kb. We compared this assembly to the reference and found that GraphUnzip did not introduce any phasing error (Figure 2).

### 3.2 Adineta vaga

We then tested GraphUnzip on the diploid genome of the bdelloid rotifer *Adineta vaga*. As this genome has variable levels of heterozygosity along its chromosomes, which include highly heterozygous regions, it has proven difficult to collapse its haplotypes into a haploid assembly [1]. High-accuracy short reads (Illumina), low-accuracy long reads (ONT and PacBio CLR) and Hi-C reads were already available [16], and we additionally sequenced high-accuracy long reads (HiFi) in order to test different assembly strategies. To improve the accuracy of ONT and PacBio CLR, they were corrected with Illumina reads and the tool Ratatosk [17]; the corrected datasets are dubbed cONT and cPacBio in what follows.

The HiFi reads were assembled using Flye and hifiasm; for corrected long reads, we only present assemblies with Flye, as hifiasm did not handle well these corrected reads (data not shown). GraphUnzip was run on these draft assemblies using long reads and/or Hi-C data. We evaluated the resulting phased assemblies based on their total size, NG50, complete single-copy and duplicated BUSCO features [21], and contact maps of the longest supercontigs (Table 1, Figure 3). All assemblies phased by GraphUnzip increased in assembly size as collapsed homozygous regions became duplicated. The NG50 of the HiFi assemblies rose from 4.8 to 11.9 Mb after GraphUnzip (using only Hi-C reads). Assemblies of cONT also reached high NG50 values when phasing with ONT/cONT and/or Hi-C (up to 16.0 Mb using both). The NG50s of cPacBio assemblies were not as high as for cONT assemblies, yet they reached a NG50 up to 4.6 Mb, whereas the initial NG50 was only 249 kb. As expected, the assemblies showed an increase in the number of complete duplicated BUSCO features. The contact maps did not display any error.

**Table 1:**
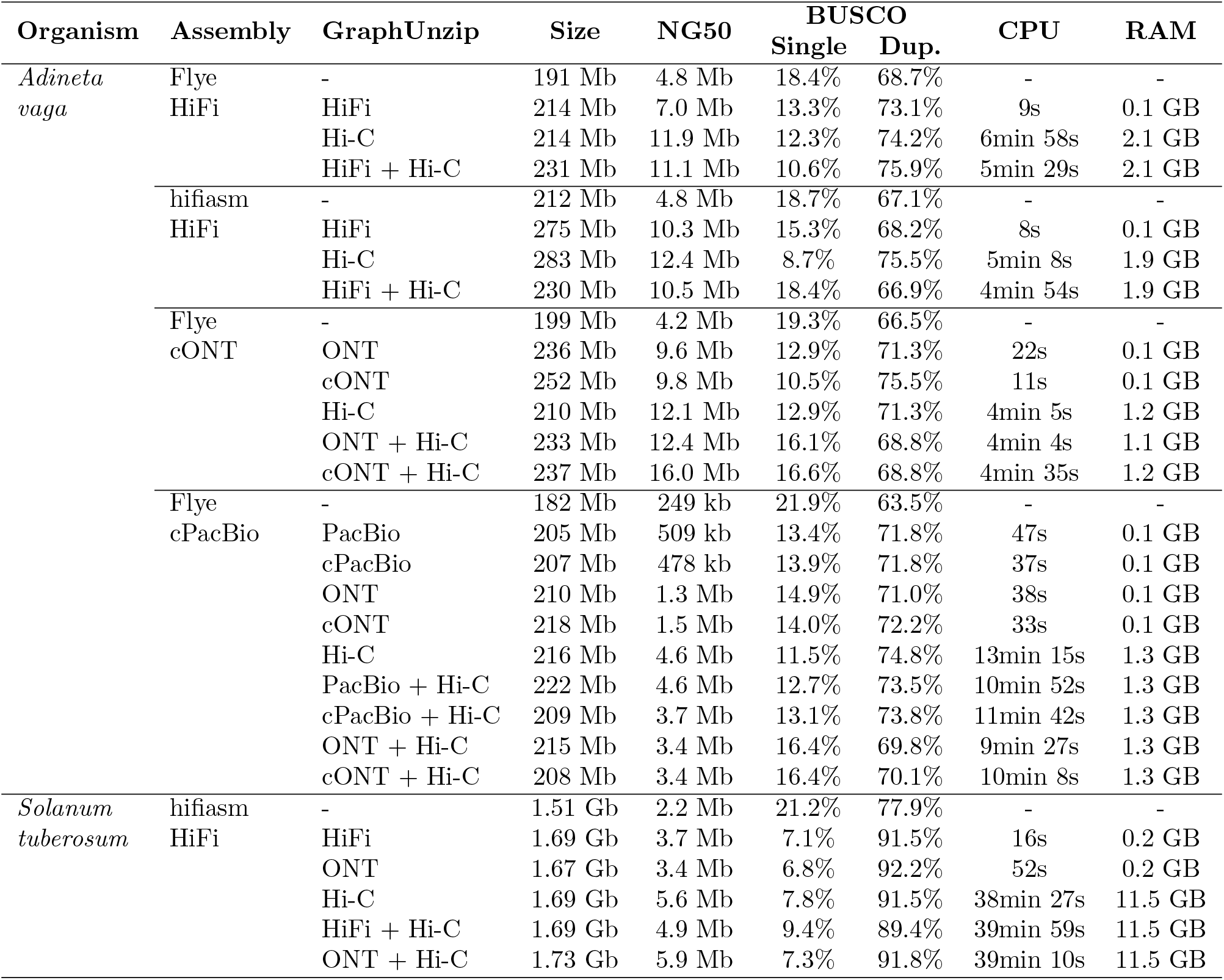
Assembly metrics. The NG50 values were computed based on an estimated genome size of 192.6 Mb for *Adineta vaga*, and 1.67 Gb for *Solanum tuberosum*.

**Figure 3:**
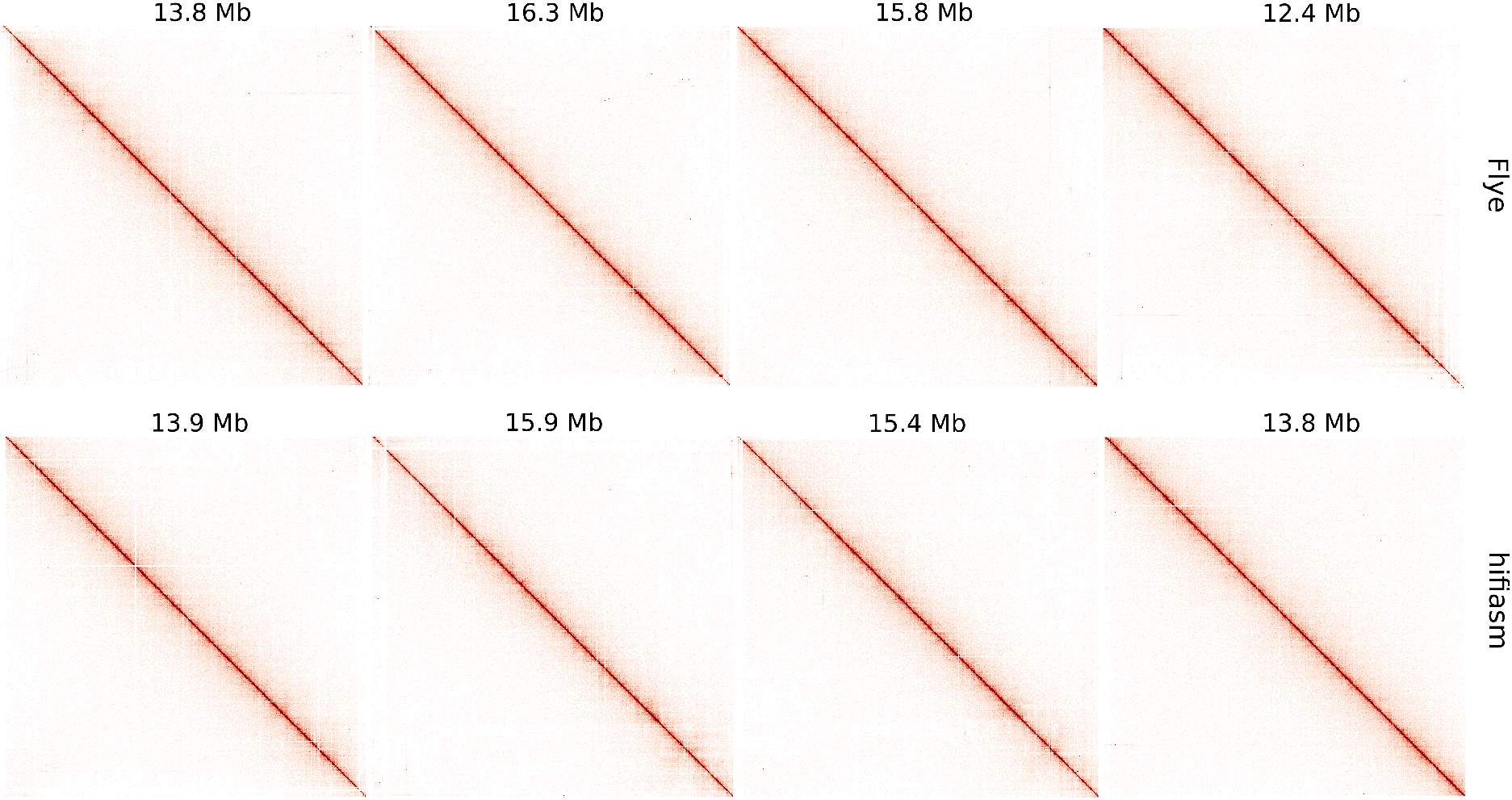
Contact maps of the four largest supercontigs after unzipping the *Adineta vaga* Flye and hifiasm assemblies of HiFi reads using GraphUnzip with HiFi and Hi-C reads.

We scaffolded the Flye and hifiasm draft assemblies of HiFi reads with ALLHiC to compare with the GraphUnzip supercontigs. For both assemblies, ALLHiC connected the contigs excessively as the scaffolds had N50s of 27.3 Mb (Flye) and 132.3 Mb (hifiasm), while the largest scaffold in the chromosome-level haploid reference is only 20.4 Mb. Besides, the ALLHiC scaffolds had poor BUSCO scores, lower than the draft assemblies: 51.8% of complete single-copy features and 35.5% of complete duplicated features for the Flye scaffolds; 74.9% and 11.5% for the hifiasm scaffolds.

### 3.3 Solanum tuberosum

To benchmark GraphUnzip against a larger assembly, we further tested it on the diploid genome of the potato *Solanum tuberosum* RH89-039-16, for which a phased assembly of 1.67 Gb [20] was recently published. We assembled the HiFi reads with hifiasm and then ran GraphUnzip using the HiFi, ONT and/or Hi-C reads. The draft assembly was 1.51 Gb, and after phasing with GraphUnzip, the assembly size rose to 1.67-1.73 Gb. GraphUnzip also increased the continuity: from 2.2 Mb, the NG50 reached 3.4 to 5.9 Mb. The combination of both ONT and Hi-C reads yielded the highest NG50. As observed with *Adineta vaga* assemblies, Hi-C reads improved the continuity better than long reads. The number of duplicated BUSCO features also increased, from 77.9% (raw assembly) to 89.4-92.2%. It should be noted that the reference sequence had only 76.9% of duplicated BUSCO features. In addition, the overall BUSCO completeness of the GraphUnzip supercontigs is slightly higher than the reference: 98.6-99.3% against 98.5% for the reference.

We also tried an assembly of the HiFi reads with Flye, but the draft assembly was only 827 Mb, little below half the expected size, which indicates that the haplotypes were collapsed. A good candidate assembly for GraphUnzip should have uncollapsed heterozygous regions, as GraphUnzip is not able to retrieve a missing haplotype in collapsed heterozygous regions and can only duplicate the remaining haplotype, leading in that case to a suboptimal result.

### 3.4 Computational performance

For both the small *Adineta vaga* genome (192.6 Mb) and the larger *Solanum tuberosum* genome (1.67 Gb), GraphUnzip required limited computational resources. For *Adineta vaga*, GraphUnzip ran in less than 15 minutes, and in less than a minute when using only long reads. The RAM usage reached a maximum of 2.1 GB. For *Solanum tuberosum*, GraphUnzip ran in less than 1 hour, and using up to 11.5 GB. The run time was also shorter when using only long reads, below 1 minute. The longer run time when using Hi-C reads was due to the building of the interaction matrix. As this interaction matrix is outputted by the program, this file can be reused for other runs, that will finish faster. Therefore, users can try several sets of parameters to optimize the result, with short runtimes.

## 4 Conclusion

GraphUnzip is a flexible tool that can phase assemblies of high-accuracy long reads (HiFi or corrected ONT or corrected PacBio CLR) with long reads and/or Hi-C. As genome projects now usually include long reads and Hi-C to obtain chromosome-level assemblies, GraphUnzip can easily be integrated in assembly projects to obtain *de novo* phased assemblies for non-model organisms.

## Author contributions statement

R.F., N.G. and J.-F.F. jointly designed GraphUnzip. R.F. and N.G. implemented GraphUnzip. R.F. and N.G. tested GraphUnzip. R.F., N.G. and J.-F.F. wrote the manuscript.

## Acknowledgments

This project was funded by the Horizon 2020 research and innovation program of the European Union under the Marie Sklodowska-Curie grant agreement No 764840 (ITN IGNITE, www.itn-ignite.eu). Part of this analysis was performed on computing clusters of the Leibniz-Rechenzentrum (LRZ) and the Consortium des Equipements de Calcul Intensif (CÉCI) funded by the Fonds de la Recherche Scientifique de Belgique (F.R.S.-FNRS) under Grant No. 2.5020.11.

